# A complex two-component system CdgKA-CdgSHA fine-tunes c-di-GMP homeostasis in a Cyanobacterium

**DOI:** 10.64898/2026.09.11.750938

**Authors:** Qing-Xue Sun, Zhao-Ru Ding, Cheng-Cai Zhang, Xiaoli Zeng

**Affiliations:** Key Laboratory of Algal Biology, Institute of Hydrobiology, Chinese Academy of Sciences, Wuhan, Hubei 430072, People’s Republic of China; Hubei Hongshan Laboratory, Wuhan 430070, People’s Republic of China; State Key Laboratory of Lake and Watershed Science for Water Security, Chinese Academy of Sciences, Nanjing 210008, People’s Republic of China; University of Chinese Academy of Sciences, Beijing 100049, People’s Republic of China

**Keywords:** Cell size, c-di-GMP, complex two-component system, cyanobacteria

## Abstract

The second messenger c-di-GMP plays a critical role in regulating cell size and growth in the cyanobacterium *Anabaena* sp. PCC 7120. While all 16 c-di-GMP metabolic enzymes and the primary receptor CdgR have been identified, how metabolic enzyme activity is controlled to maintain c-di-GMP homeostasis remains largely unexplored. Here, using a suppressor-screening approach based on c-di-GMP-deficient mutants, we identify All0729 (CdgKA) as a regulator that orchestrates c-di-GMP homeostasis and cell size regulation. Genetic and quantitative analyses demonstrate that CdgKA acts upstream of All2306 (CdgSHA), the reported major c-di-GMP hydrolase, to modulate the activity of the latter. While c-di-GMP synthesis mutants displayed reduced c-di-GMP levels and decreased cell size, inactivation of *cdgKA* or *cdgSHA* in these mutant backgrounds rescued both the c-di-GMP levels and cell size. We propose that CdgKA and CdgSHA form a complex two-component system, and the phosphorylation of CdgSHA enhances its c-di-GMP-degrading activity. While the previously identified two-component system composed of CdgK-CdgS regulates c-di-GMP synthesis, the CdgKA-CdgSHA system controls its degradation. Together, they constitute a dual TCS network that precisely tunes c-di-GMP levels and cell size, establishing a regulatory link between two-component signaling and second messenger homeostasis in cyanobacteria.

**Importance:** Cyanobacteria are photosynthetic microbes being developed for sustainable fuel and chemical production, but their growth efficiency depends on maintaining an appropriate cell size. This study identifies a molecular pathway with CdgKA-CdgSHA in a cyanobacterium that controls cell size and growth by the key signaling molecule c-di-GMP. This pathway controls c-di-GMP degradation, while a previously identified CdgK-CdgS signaling system controls c-di-GMP synthesis. These systems together fine-tune c-di-GMP homeostasis in response to environmental conditions. This work provides a foundation for engineering cyanobacteria with improved growth characteristics for industrial applications.

## Introduction

In prokaryotes, two-component systems (TCSs) constitute a major signal transduction strategy that couples environmental or cellular stimuli to adaptive physiological responses, including transcriptional reprogramming, behavioral modulation, proteolysis, and differentiation (1–4). A canonical TCS comprises a sensor histidine kinase (HK) and a response regulator (RR). The HK usually contains sensor domains, a kinase domain, and a histidine phosphotransfer (HPT) domain, while the RR consists of phosphoacceptor receiver (REC) domains and effector domains (4–6). Upon signal perception, the HK auto-phosphorylates and transfers the phosphoryl group to its cognate RR, which in turn regulates gene expression, enzyme activity, RNA-protein interaction, or protein-protein interaction (4, 5). In many cases, intermediary Hpt, HK, and REC domains participate in multistep phosphorelays, thereby expanding regulatory complexity (1, 2).

c-di-GMP is a ubiquitous bacterial signaling molecule that plays a crucial regulatory role in various cellular processes, such as motility, adhesion, biofilm formation, development, morphogenesis, and virulence (7–10). Its intracellular concentration is dynamically balanced by the opposing activities of diguanylate cyclases (DGCs), which harbor GGDEF domains for synthesis, and phosphodiesterases (PDEs), which carry EAL or HD-GYP domains for degradation (8, 9, 11). To achieve precise spatiotemporal control, bacteria typically encode a diverse array of DGCs and PDEs (12–15). For example, *E. coli* and *Pseudomonas aeruginosa* contain 29 and 41 such enzymes, respectively (13, 16). These enzymes are often modular proteins, with the GGDEF, EAL, or HD-GYP domains coupled with various sensory domains located in their N-terminals, including the Per-ARNT-Sim (PAS) domain, the GAF domain, the HAMP domain, and the REC domain, facilitating a rapid and precise modulation of intracellular c-di-GMP levels in response to various environmental and cellular signals (8, 9, 11). c-di-GMP regulates bacterial physiological processes and environmental adaptation through its downstream receptors (9, 10, 12, 14, 17). This intricate interplay forms a complex c-di-GMP regulatory network, contributing to the accurate regulation of bacterial physiological responses (12, 16, 18).

*Anabaena* PCC 7120 (hereafter *Anabaena*) is a filamentous cyanobacterium that contains 16 genes encoding proteins for c-di-GMP metabolism (15, 19–22). Our recent work established that c-di-GMP not only modulates cell size but is also indispensable for cell viability in *Anabaena* (17, 21). Through systematic analysis, we identified the major DGCs and PDEs governing c-di-GMP homeostasis and defined two critical physiological thresholds: a moderate reduction (∼80%±5% of wild-type levels, morphological threshold) impairs cell size, whereas a severe drop (below ∼50%±5%, viability threshold) becomes lethal. The 16 enzymes operate in a manner analogous to an electromechanical dual-relay system: thirteen enzymes constitute a baseline signaling module, the c-di-GMP synthases CdgS and All1219 act as a responsive relay, and Alr3599 serves as an emergency relay activated only when c-di-GMP drops to lethal levels. Meanwhile, Alr2306, a protein containing both GGDEF and EAL domains, was identified as one of the major PDEs responsible for maintaining intracellular c-di-GMP levels (21). Furthermore, both cell size and viability effects of c-di-GMP are mediated by the receptor CdgR, which modulates cell size partly through interaction with the global transcriptional factor DevH (17, 23). Despite these advances, the upstream regulatory mechanisms that control the activities of these c-di-GMP metabolic enzymes remain largely unexplored.

In this study, we identified a histidine kinase, All0729 (termed CdgKA for <u>c</u>-<u>d</u>i-<u>G</u>MP two-component signaling <u>k</u>inase <u>A</u>), through suppressor screening by the reported cell growth-defective mutant *Δ8DGC-1*. Extensive genetic evidence demonstrates that CdgKA and Alr2306 (termed CdgSHA for <u>c</u>-<u>d</u>i-<u>G</u>MP <u>s</u>ynthase and <u>h</u>ydrolase <u>A</u>) constitute a complex TCS, where CdgKA-mediated phosphorylation promotes the PDE activity of CdgSHA, thereby promoting c-di-GMP degradation. We further show that the CdgKA-CdgSHA system regulates cell size and growth in *Anabaena*. Together with the previously characterized CdgK-CdgS system (22), which controls c-di-GMP synthesis, our findings reveal that cell-size maintenance by c-di-GMP requires the coordination of two systems.

## Results

### Inactivation of *cdgKA* restores the phenotype of cell size reduction and loss of viability in the *8ΔDGC-1* mutant strain

To explore the upstream regulatory mechanisms of the c-di-GMP-dependent cell size control pathway, we performed a suppressor screen using the growth-defective 8*ΔDGC-1* strain, in which all 8 genes encoding enzymes in DGC-only form were deleted (*ΔcdgSΔalr3504Δalr3599Δall2416Δall1012Δall5174Δall4896CT-all1219*) (21). As described previously, the expression of *all1219* is controlled by the CT promoter in order to maintain the viability of the mutant by supplying inducers in the culture medium (21). The *8ΔDGC-1* strain fails to grow in BG11 lacking Cu²⁺ and TP due to its intracellular c-di-GMP level dropping below 50%±5% of the WT level (Figure 1A and 1B) (21). From this background, we isolated 100 suppressor mutants that regained growth under the non-permissive conditions (Figure 1).

**Figure 1.**
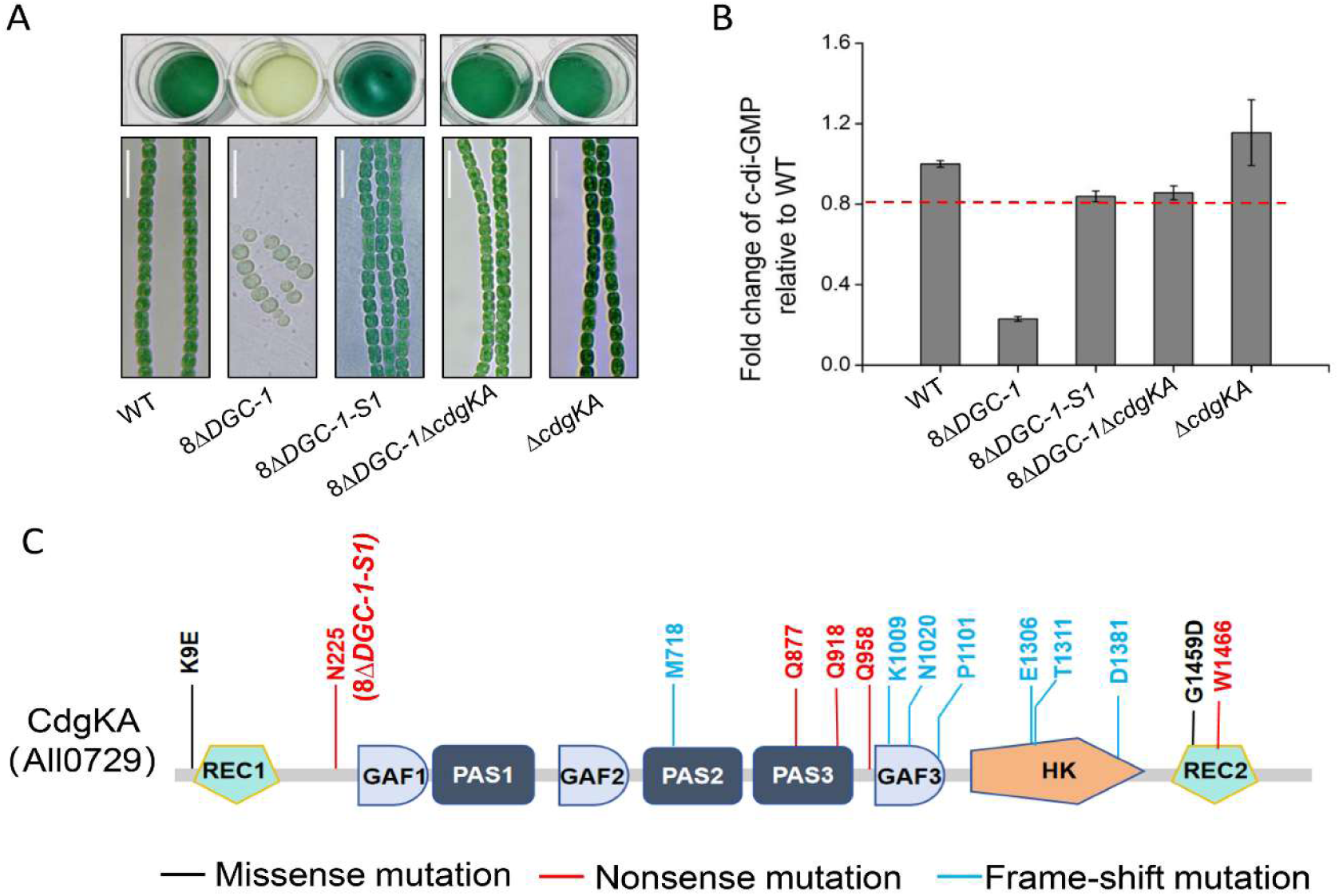
Inactivation of *cdgKA* suppresses the phenotype of the 8*ΔDGC-1* mutant. (A) The growth (top panel) and micrographs (Bottom panel) of indicated strains cultured in BG11 medium without Cu^2+^ and theophylline. The growth of different strains was tested in 24-well plates. All cultures started with a similar OD at 0.3 diluted from a pre-culture, and were imaged after 4 days of incubation. *8ΔDGC-1-S1*: Suppressor mutation of *8ΔDGC-1*. Scale bars, 15 μm. (B) The intracellular c-di-GMP levels of the indicated strains at 96 h in BG11 medium without Cu^2+^ and theophylline. The c-di-GMP concentrations of all mutants were normalized to the levels of the WT (wild-type) strain. The fold change numbers are shown as mean ± SD from three biological replicates. Red dotted line represents 80% of the WT c-di-GMP level. 8*ΔDGC-1: ΔcdgSΔalr3504Δalr3599Δall2416Δall1012Δall5174Δall4896 CT-all1219*. (C) The schematic representation of frameshift, missense, and nonsense mutations in CdgKA of the *8ΔDGC-1* suppressors. Protein domains were analyzed using SMART (http://smart.embl-heidelberg.de/).

Whole-genome sequencing of selected suppressors revealed mutations in two genes: *cdgR* and *cdgKA*. CdgR is a known downstream c-di-GMP receptor that transduces the c-di-GMP signal, and its deletion could rescue the cell size reduction phenotype caused by decreased c-di-GMP levels (17). We further amplified and sequenced the *cdgKA* coding region of all the obtained suppressors. The results demonstrated that mutations occurring within *cdgKA* represented 30% of all suppressor mutations, corresponding to 14 distinct mutation sites, including 7 frameshift mutations and 7 missense or nonsense mutations (Figure 1C). One representative suppressor, *8ΔDGC-1-S1*, which carries an N225 nonsense mutation in *cdgKA*, was subjected to phenotype analysis. The *8ΔDGC-1-S1* strain regained growth ability comparable to WT in BG11 lacking Cu²⁺ and TP. Microscopic observation further revealed that its cell size was restored to WT levels (Figure 1A). In contrast, the *8ΔDGC-1* strain exhibited severe cell size reduction and failed to grow, consistent with a previous report (21). Collectively, these results suggest that inactivation of *cdgKA* effectively rescues the growth arrest and cell size defects of the *8ΔDGC-1* mutant.

To further confirm that *cdgKA* inactivation is responsible for the suppressor phenotype of *8ΔDGC-1*, we constructed a markerless in-frame deletion of *cdgKA* in the *8ΔDGC-1* background (*8ΔDGC-1ΔcdgKA*). A single *ΔcdgKA* strain was also generated as a control. As shown in Figure 1A, the *8ΔDGC-1ΔcdgKA* exhibited growth and cell size comparable to the WT, completely abolishing the growth arrest and cell size reduction observed in the *8ΔDGC-1* strain. The *ΔcdgKA* single mutant displayed no particular phenotype under the conditions tested. Taken together, these results confirm that *cdgKA* inactivation dominantly rescues the *8ΔDGC-1* phenotypes, placing *cdgKA* within the c-di-GMP regulation pathway.

### *cdgKA* regulates cell size and cell growth by modulating c-di-GMP levels

To determine the role of *cdgKA* in the c-di-GMP signaling pathway, we measured intracellular c-di-GMP levels in the five strains: *8ΔDGC-1-S1*, *8ΔDGC-1ΔcdgKA*, *ΔcdgKA,* and the WT and *8ΔDGC-1* as a control. As shown in Figure 1B, the c-di-GMP level in *8ΔDGC-1-S1* increased to approximately 84%±2.7% of the WT level, whereas the *8ΔDGC-1* strain showed a severe reduction. Similarly, *8ΔDGC-1ΔcdgKA* exhibited a c-di-GMP concentration that is about 86%±3.4% of the WT level. Thus, in both the suppressor mutant (*8ΔDGC-1-S1*) and the reconstructed mutant (*8ΔDGC-1ΔcdgKA*), the c-di-GMP levels were above the threshold (∼80%±5%) required for cell size maintenance (21). *ΔcdgKA* showed an approximately 1.2±0.16-fold increase in c-di-GMP levels relative to WT (Figure 1B), consistent with the previous finding showing that an increase in c-di-GMP in *Anabaena* causes no particular phenotype (20, 21). These results indicate that *cdgKA* inactivation rescues the *8ΔDGC-1* phenotypes by upregulating c-di-GMP levels.

To further confirm that *cdgKA* indeed regulates the intracellular c-di-GMP levels in *Anabaena,* we constructed a *ΔcdgSΔcdgKA* double mutant, as CdgS has been previously identified as the primary DGC involved in cell size regulation (15, 21, 22). Deletion of *cdgKA* suppressed the reductions in both cell length and width caused by *cdgS* deletion (Figure 2A and 2B). Consistently, the intracellular c-di-GMP level of the *ΔcdgSΔcdgKA* mutant was restored to approximately 1.3±0.05-fold of the WT level (Figure 2C). As a control, the complemented strain *ΔcdgSΔcdgKA::cdgKA* reverted to the cell size and c-di-GMP level observed in the *ΔcdgS* mutant (Figure 2). Together, these results demonstrate that CdgKA modulates intracellular c-di-GMP levels to regulate cell size and growth in *Anabaena*, and indicate that it functions upstream of the c-di-GMP regulation pathway.

**Figure 2.**
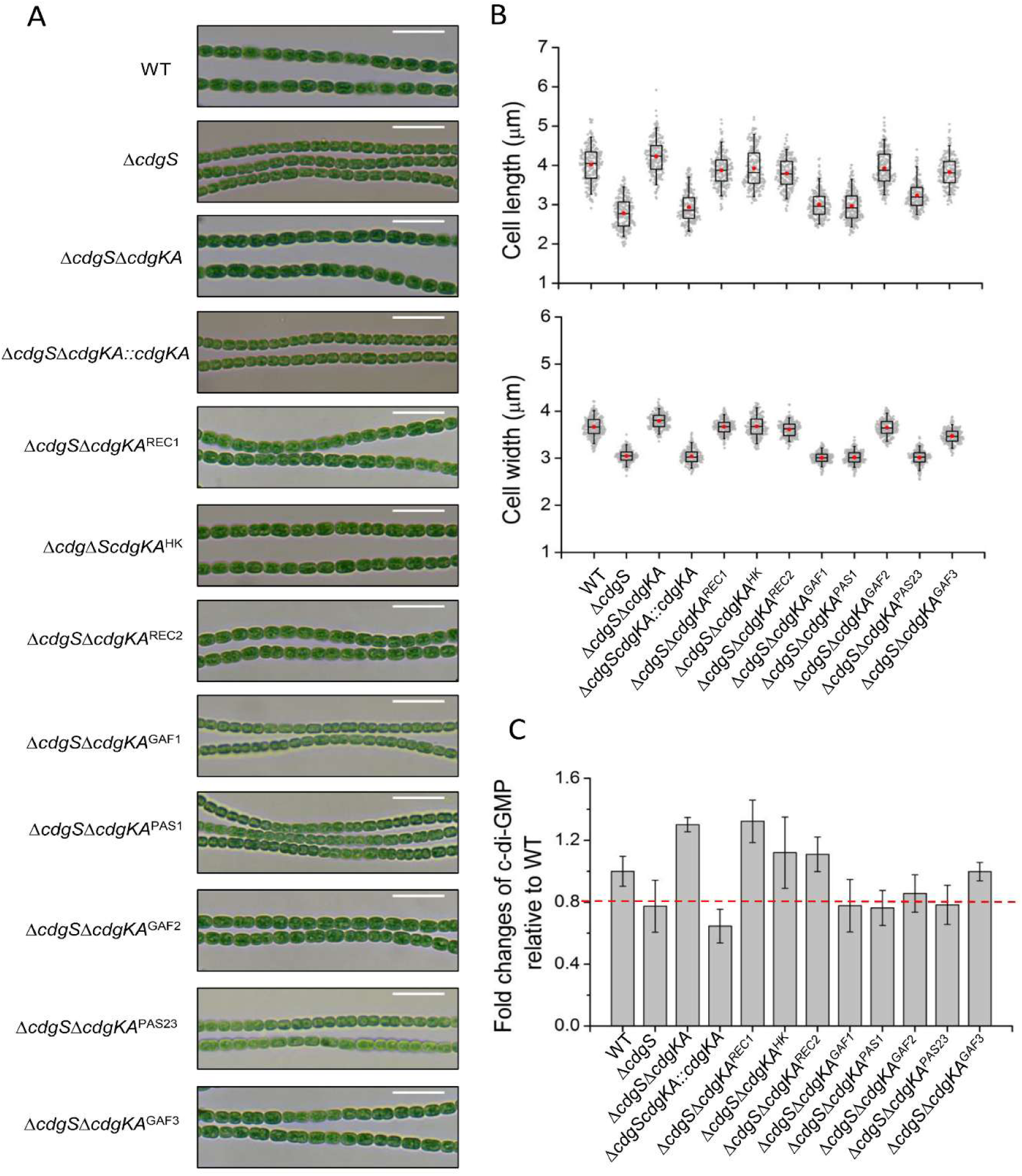
The phenotype of different truncated mutants of *cdgKA* in the background of *ΔcdgS*. (A) Micrographs of indicated strains cultured in BG11 medium. Scale bars: 15µm. (B) Cell size parameters (cell length and width) of the indicated strains. The cell length and cell width were measured based on images, as shown in (A). 200 cells of each strain from three independent experiments were measured. The boxplots enclose the 25th and the 75th percentile, with the black line representing the median value and the red dot representing the mean value. (C) Intracellular c-di-GMP levels of the indicated strains at 96 h in BG11 medium. The c-di-GMP concentrations of all mutants were normalized to the levels of the wild-type strain. The fold change numbers are shown as mean ± SD from three biological replicates.

### Functional dissection of the histidine kinase CdgKA domains involved in c-di-GMP level control

CdgKA was annotated as a sensor histidine kinase, featuring an N-terminal REC domain (REC1), three GAF domains (GAF1, GAF2, and GAF3) and three PAS domains (PAS1, PAS2, and PAS3) distributed in the middle region, and an HK and a REC domain (REC2) at the C-terminal region (Figure 1C). To investigate the functional contribution of individual domains, we constructed a series of in-frame deletion mutants in the *ΔcdgS* background and analyzed their cell size and intracellular c-di-GMP levels (Figure 2 and Figure S1). We first generated three truncated mutants—*ΔcdgSΔcdgKA^REC1^*, *ΔcdgSΔcdgKA^HK^*, and *ΔcdgS ΔcdgKA^REC2^* (Figure 2 and S1)—which lack the REC1, HK, and REC2 domains of CdgKA in the *ΔcdgS* background, respectively. As shown in Figure 2, all three mutants restored cell length and width to those of WT and *ΔcdgSΔcdgKA* double mutant (Figure 2A and 2B), with c-di-GMP levels reaching approximately 1.2-fold of the WT level, comparable to those of the *ΔcdgSΔcdgKA* double mutant (Fig. 2C). In contrast, the *ΔcdgS* mutant exhibited reduced cell size and a c-di-GMP level lower than 80%±5% of the WT level (Figure 2) (21, 22). These results indicate that REC1, HK, and REC2 domains are each required for CdgKA function, suggesting that CdgKA acts as a histidine kinase that controls cell size through phosphorylation signaling.

We constructed five additional mutants with deletions of individual predicted sensor domains of CdgKA in the *ΔcdgS* background: *ΔcdgSΔcdgKA^GAF1^*, *ΔcdgSΔcdgKA^PAS1^*, *ΔcdgSΔcdgKA^GAF2^*, *ΔcdgSΔcdgKA^PAS23^* (PAS2 and PAS3 deleted together), and *ΔcdgSΔcdgKA^GAF3^* (Figure 2 and S1). The *ΔcdgSΔcdgKA^GAF1^*, *ΔcdgSΔcdgKA^PAS1^*, *and ΔcdgSΔcdgKA^PAS23^* mutants displayed cell size defects and reduced c-di-GMP levels similar to those of the *ΔcdgS* strain, indicating that deletion of these domains failed to suppress the phenotype of *ΔcdgS,* unlike the effect of *cdgKA* inactivation (Figure 2). In contrast, the *ΔcdgSΔcdgKA^GAF2^* and *ΔcdgSΔcdgKA^GAF3^* mutants exhibited cell sizes comparable to that of the WT and c-di-GMP levels above 80%±5% of the WT level, resembling the *ΔcdgSΔcdgKA* phenotype. Collectively, these results suggest that GAF2 and GAF3 function as the critical sensor domains for the activity of CdgKA, whereas GAF1, PAS1, and PAS23 are dispensable for cell size regulation under the tested conditions.

### CdgKA functions in the same genetic pathway as the major PDE CdgSHA to regulate c-di-GMP levels

Since CdgKA lacks DGC or PDE domains, it likely regulates c-di-GMP levels by modulating the activity of one or more of the remaining eight c-di-GMP turnover enzymes in *8ΔDGC-1*. The eight candidate enzymes include six dual-function DGC-PDEs (All0219, All1175, CdgSHA, Alr3170, All4225, All4897) and two PDEs (Alr1230, Alr3920) (21). Among them, only CdgSHA, Alr1230, and Alr3920 possess the REC domain (Figure 3A), making them potential targets of the histidine kinase CdgKA. Our previous work identified CdgSHA as the major PDE in *Anabaena* (21). Among all c-di-GMP metabolic enzymes, CdgSHA is the only one whose deletion elevates intracellular c-di-GMP levels in WT *Anabaena* and partially compensates for the reduced c-di-GMP levels and cell size of the *ΔcdgS* mutant (21). This phenotype is similar to that observed in the *ΔcdgSΔcdgKA* strain (Figure 2). These findings suggest that CdgKA may modulate the activity of CdgSHA, thereby regulating c-di-GMP levels and consequently affecting cell size and growth.

**Figure 3.**
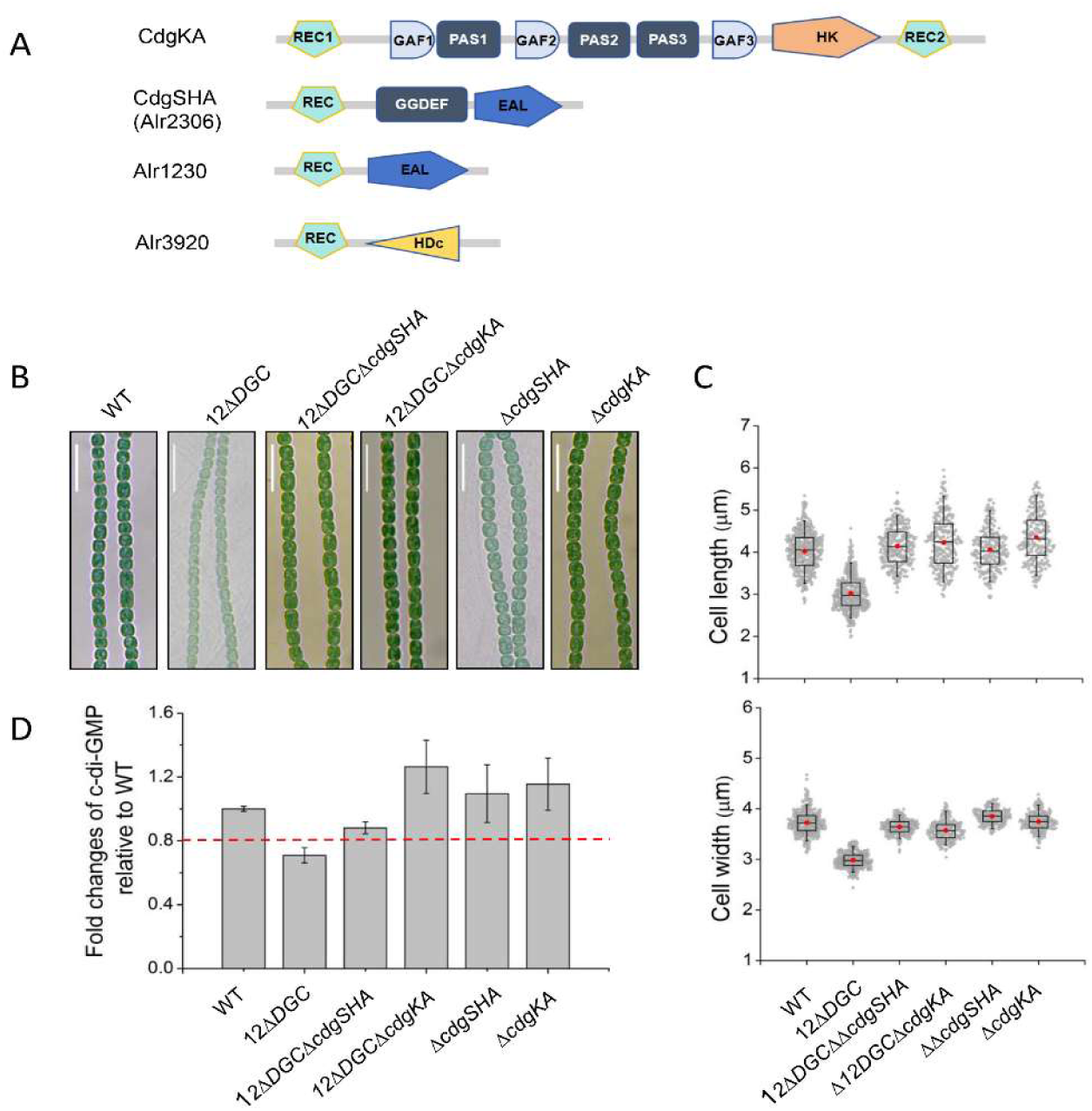
CdgKA functions in the same genetic pathway as CdgSHA. (A) Predicted domain architectures of indicated proteins. Protein domains were analyzed using SMART (http://smart.embl-heidelberg.de/). (B) Micrographs of indicated strains cultured in BG11 medium without Cu^2+^ and theophylline. Scale bars: 15 μm. (C-D) The cell size parameters (cell length and width) and intracellular c-di-GMP levels of the indicated strains as shown in (B). Red dotted line in (D) represents 80% of the WT c-di-GMP level. 12*ΔDGC*: *ΔcdgSΔalr3504Δalr3599Δall2416 Δall1012Δall4896Δall4897Δalr3170Δall0219Δall4225Δall1175Δall5174*.

To further confirm this hypothesis, we constructed a 1*2ΔDGCΔcdgKA* mutant, in which *cdgKA* was deleted in the *12ΔDGC* background. This background was chosen over 8ΔDGC because it retains only four c-di-GMP metabolic enzymes: the three putative REC-domain-containing targets of CdgKA (CdgSHA, Alr1230, and Alr3920) and the major DGC All1219 (21), thereby minimizing genetic redundancy and enabling a direct assessment of the role of CdgSHA in CdgKA-mediated regulation of c-di-GMP levels and cell size. *12ΔDGC* displayed cell size defects and significantly reduced c-di-GMP levels, as reported previously (21). Further analysis revealed that *12ΔDGCΔcdgKA* restored cell size to that of the *12ΔDGCΔcdgSHA* (also named *13ΔDGC* in the previous report) strain, with both mutants recovering to WT levels (Figure 3B and 3C). Consistently, intracellular c-di-GMP levels in both strains were restored to above 80% of the WT level (Figure 3D). These results indicate that inactivation of either *cdgKA* or *cdgSHA* produces similar phenotypes in the *12ΔDGC* background, supporting that CdgKA regulates cell size through modulating CdgSHA activity. Quantitative measurements revealed a subtle but consistent difference in the function of the two genes: the c-di-GMP level of *12ΔDGCΔcdgKA* reached approximately 1.3±0.17-fold of the WT level, whereas that of *12ΔDGCΔcdgSHA* was approximately 88%±3.7% of WT (Figure 3D). This distinction suggests that CdgSHA, which harbors both GGDEF and EAL domains, may function as a dual-functional enzyme whose DGC activity also contributes to the intracellular c-di-GMP pool for cell growth, in addition to its PDE activity potentially regulated by CdgKA.

### Phosphorylation signal within the CdgKA and CdgSHA is important for c-di-GMP homeostasis

The truncation experiments described above showed that deletion of any of the three phosphorelay domains—REC1, HK, or REC2—abolished CdgKA function (Figures 2 and 3). To determine whether this functional requirement reflects phosphotransfer activities, we replaced each of the conserved phosphoacceptor residues within each domain, H1147 in HK, D57 in REC1, and D1477 in REC2, with alanine in the *ΔcdgS* background (Figure 4A). The *ΔcdgSΔcdgKA^H1147A^* and *ΔcdgSΔcdgKA^D1477A^* mutants both exhibited cell sizes comparable to WT with a concomitant recovery of the c-di-GMP levels up to 80% of the WT level, whereas *ΔcdgSΔcdgKA^D57A^* showed *a* similar phenotype as the *ΔcdgS* single mutant (Figure 4B–4C). These results suggest that H1147 and D1477 are involved in autophosphorylation and phosphotransfer expected for a histidine kinase, consistent with the data obtained with domain truncations (Figure 2). In contrast, the phosphorylation site D57 in REC1 is dispensable for CdgKA function under the test conditions, implying that the REC1 domain may serve a sensor domain rather than a phosphorelay role, whereas its truncation disrupts the activity of CdgKA.

**Figure 4.**
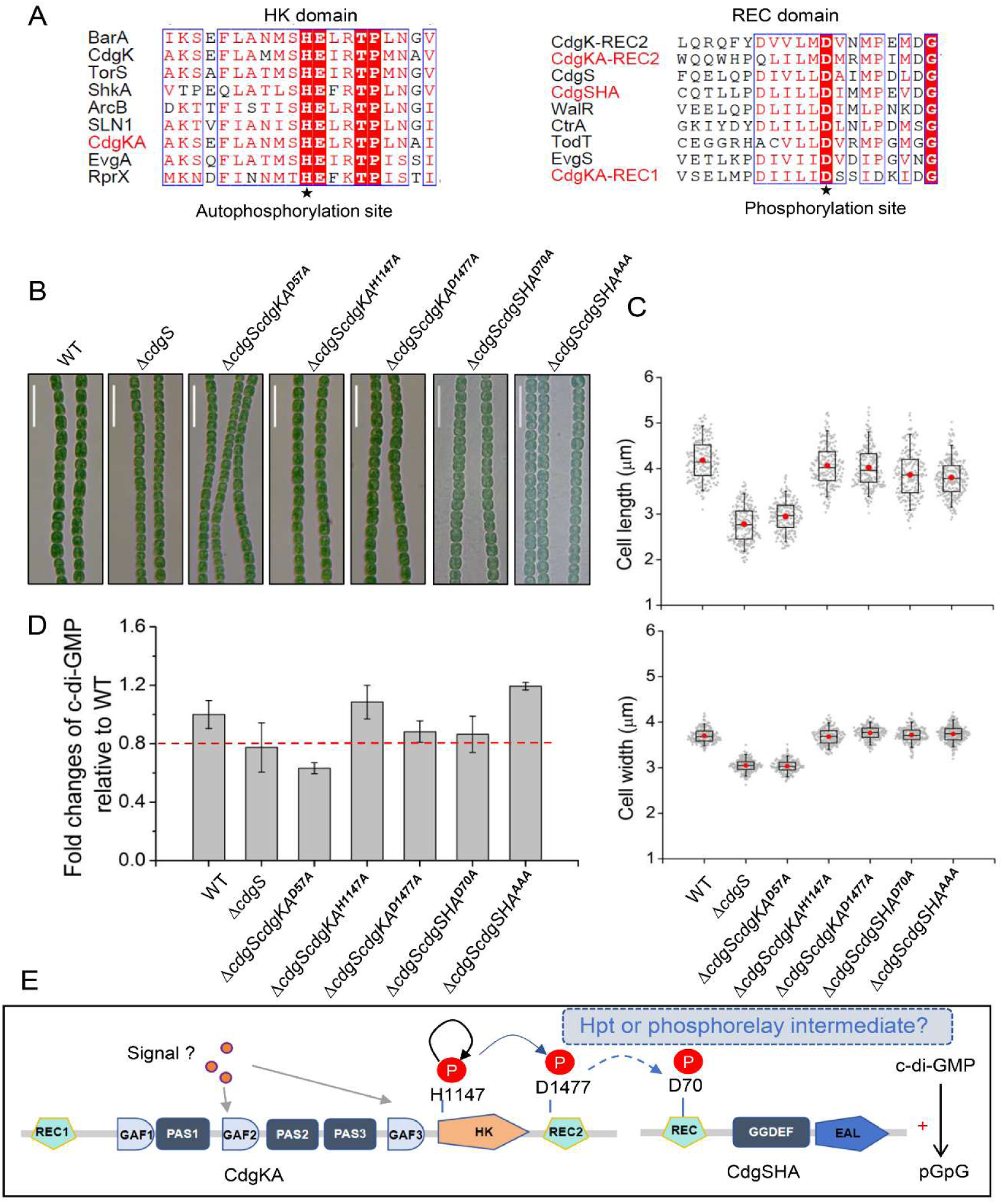
CdgKA-CdgSHA forms a complex two-component system in c-di-GMP control. (A) Sequence alignment of the HK and REC domains from different TCS systems. All homologous domains of proteins were retrieved from NCBI (https://www.ncbi.nlm.nih.gov/). BarA (Genebank: APC52989.1); TorS (GenBank: AAF96608.1); EvgA (Genebank: BAA16241.1); ArcB (Genebank: CAA37397.1); LSN1 (Genebank: AJR37544.1); RprX (Genebank: AAB26173.1); WalR (Genebank: NP_391921); CdgK (Genebank: WP_010997026.1); CdgS (Genebank: WP_010997025.1); CtrA (Genebank: WP_010920871.1); TodT (Genebank: APC70898.1); EvgS (Genebank: BAA16240.1). (B) Micrographs of indicated strains cultured in BG11 medium. Scale bars, 15 μm. (C-D) The cell size parameters (cell length and width) and intracellular c-di-GMP levels of the indicated strains as shown in (B). The red dotted line in (D) represents 80% of the WT c-di-GMP level. Length and width of at least 200 cells of each strain in (B) were measured. The c-di-GMP concentrations of all mutants in (D) were normalized to the levels of the wild-type strain. The fold change numbers are shown as mean ± SD from three biological replicates. (E) The schematic representation of signal transduction from CdgKA to CdgSHA. The environmental signal was preserved by the GAF2 and GAF3 domains of CdgKA under the tested conditions. The unknown Histidine Phosphotransferase (Hpt) or other phosphorelay intermediates are a bridge to transduce the phosphorylation signal from CdgKA to CdgSHA. P: phosphoryl group.

CdgSHA contains an N-terminal REC domain with a conserved aspartate residue (D70), which may serve as the phosphoryl acceptor from its cognate HK (Figures 3A and 4A). To test this, we constructed a mutant, *ΔcdgSΔCdgSHA^D70A^*, in which D70 of CdgSHA was changed to Ala in the *ΔcdgS* genetic background. Phenotypic analysis revealed that this mutant exhibited cell size comparable to that of WT (Figure 4B–4C), and its intracellular c-di-GMP level (∼86.4%±12.4% of WT level) exceeded the morphological threshold (∼80%±5%) required for cell size maintenance, in contrast to the *ΔcdgS* single mutant (∼77.4%±16.8%) (Figures 2C and 4D). These results suggest that D70 is critical for CdgSHA function. As a control, we also analyzed the *ΔcdgSΔcdgSHA^AAA^* mutant, in which the conserved EAL motif required for PDE activity of CdgSHA was replaced with Ala residues. This mutant phenocopied *ΔcdgSΔcdgSHA^D70A^*, suppressing both cell size and c-di-GMP reduction observed in *ΔcdgS* (Figures 4B–4D). Taken together, these findings indicated that CdgSHA functions as a response regulator whose phosphorylation at D70 regulates its PDE activity, which is required for c-di-GMP homeostasis.

## Discussion

c-di-GMP plays a critical role in regulating cell size and growth in *Anabaena* (17, 21). We previously established that CdgS, All1219, and Alr3599 serve as the primary and emergency DGCs, while CdgSHA functions as the primary PDE among the 16 c-di-GMP metabolic enzymes in *Anabaena*, forming a core network that maintains c-di-GMP homeostasis (21). c-di-GMP controls cell size and viability with thresholds of 80%±5% and 50%±5% relative to WT, respectively (21). CdgSHA is a dual-function enzyme with dominant PDE activity, harboring GGDEF and EAL domains at its C-terminus and an N-terminal REC domain characteristic of TCS response regulators (5, 21). However, the regulatory mechanism governing CdgSHA activity has remained elusive. In this study, we demonstrate that CdgKA functions as a histidine kinase that modulates CdgSHA PDE activity through phosphorylation to maintain the intracellular c-di-GMP pool. Our genetic evidence supports a model in which CdgKA and CdgSHA constitute a complex two-component system with H1147 as an autophosphorylation site in the HK domain of CdgKA; then the phosphoryl group could be transferred to D1477 in its REC2 domain, and subsequently to D70 in CdgSHA, likely via an unidentified bridging histidine phosphotransferase (Hpt) or other phosphorelay intermediate(s) (2, 4). The phosphorylation of CdgSHA could enhance its PDE activity, leading to decreased c-di-GMP levels (Figure 4E).

Through a suppressor screen in the c-di-GMP-reduced *8ΔDGC-1* strain, we isolated 30 independent mutants with mutations in *cdgKA* that restored both growth and cell size of the initial mutant strain (Figure 1). Genetic analysis across multiple cell size-deficient backgrounds, including *8ΔDGC-1*, *ΔcdgS*, and *12ΔDGC*, confirmed that *cdgKA* inactivation rescues c-di-GMP levels (Figure 2 and 3). These results establish CdgKA as a negative regulator of c-di-GMP levels by promoting its degradation. CdgKA is a hybrid HK comprising nine predicted domains (Figure 1). Domain dissection, combined with point mutation analysis of conserved residues, identified that H1147 in the HK domain and D1477 in the REC2 domain are essential for its function (Figure 2 and 4). Within the sensor module, GAF2 and GAF3 were indispensable for CdgKA activity, whereas GAF1, PAS1, and PAS23 were not (Figure 2). These results implicate GAF2 and GAF3 as important elements in perceiving unknown signals that modulate the kinase activity of CdgKA, consistent with the known ligand-binding functions of GAF domains in other sensor kinases (24–26). The identification of the precise signals sensed by CdgKA remains a key challenge for future investigation.

Multiple lines of genetic evidence support that CdgSHA is the terminal response regulator for CdgKA and that the two proteins form a functional and complex TCS (Figures 3 and 4). First, loss of *cdgKA* or *cdgSHA* in WT or various genetic backgrounds produced similar phenotypes, including restored cell size and elevated c-di-GMP pools (Figures 2 and 3). Second, substitution of the conserved aspartate D70 in the REC domain of CdgSHA abolished its function (Figures 4A-4D), implying that phosphorylation at this residue is essential for its PDE activity. Third, since the phosphoryl group could not be transferred from the REC2 domain of CdgKA to the REC domain of CdgSHA directly, there must be an additional Hpt protein or other phosphorelay intermediate(s) between CdgKA and CdgSHA, which would make this two-component system more complex. The identification of these intermediates remains a key challenge for future investigation. It is interesting to note that in various genetic backgrounds with c-di-GMP reduction, deletion of *cdgKA* consistently resulted in higher c-di-GMP levels than deletion of *cdgSHA* (Figures 2-4), suggesting that CdgSHA, although predominantly functioning as a PDE, retains intrinsic DGC activity that contributes to c-di-GMP synthesis.

The previously characterized CdgK-CdgS TCS system regulates cell size through c-di-GMP synthesis (22). In contrast, our discovery of the CdgKA-CdgSHA complex TCS system controls c-di-GMP degradation. These results reveal that cell size maintenance in *Anabaena* requires the coordination of two TCSs with opposing functions. This dual system enables *Anabaena* to precisely balance synthesis and degradation of the key second messenger c-di-GMP in response to distinct environmental or developmental cues, ensuring that c-di-GMP levels remain at the threshold required for cell size maintenance and cell growth (Figure 5). When CdgK-CdgS is absent, degradation of c-di-GMP by the CdgKA-CdgSHA system disrupts this balance, leading to cell size reduction. In contrast, when both systems are absent, the c-di-GMP pool is rebalanced above the threshold level, so that cell size reduction is suppressed (Figure 2). These dual-TCS models provide a mechanistic framework for understanding how *Anabaena* integrates multiple signals to maintain c-di-GMP homeostasis. These findings significantly enrich the c-di-GMP signaling framework in *Anabaena* and provide a foundation for future investigations into signal perception and the molecular mechanisms underlying cell size control in cyanobacteria.

**Figure 5.**
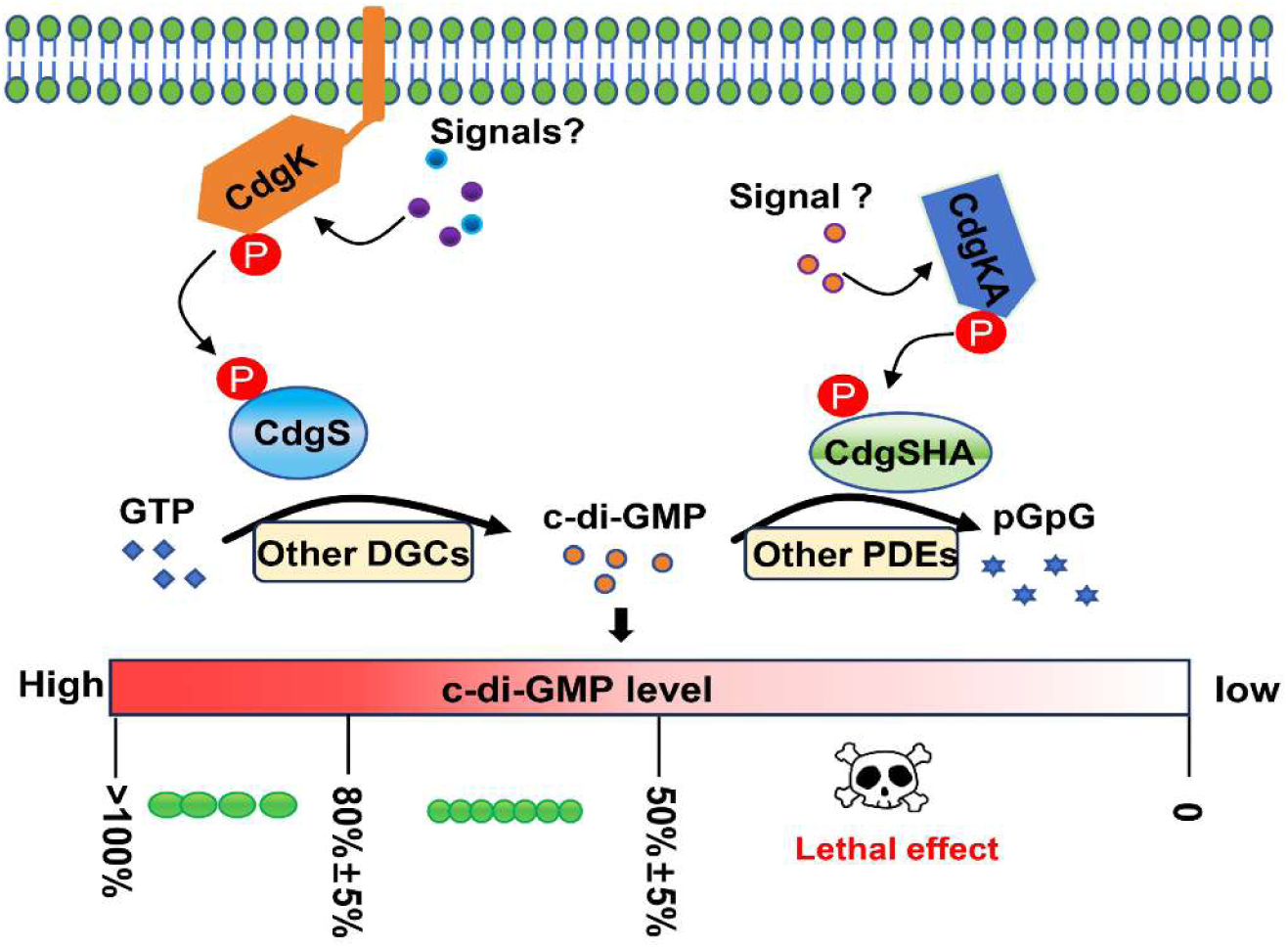
A dual TCS network that precisely tunes c-di-GMP levels to control cell fate in *Anabaena*. Sixteen c-di-GMP metabolic enzymes in *Anabaena* form a core network that maintains c-di-GMP homeostasis. c-di-GMP controls cell size and viability, with thresholds of 80%±5% and 50%±5% relative to WT, respectively. CdgS is the primary DGC, forming a TCS system with CdgK to regulate cell fate through c-di-GMP synthesis. CdgSHA functions as the primary PDE, forming a complex TCS system with CdgKA to regulate cell fate through c-di-GMP degradation. This dual system integrates diverse signals to precisely balance c-di-GMP synthesis and degradation, thereby sustaining the messenger within the permissive range for normal cell size and growth.

## Materials and methods

### Bacterial strains and growth conditions

*Anabaena* sp. strain PCC 7120 and its derivatives were cultivated in BG11 medium with shaking (30 °C, 180 rpm) and continuously illuminated with 30 μmol m^−2^ s^−1^ of white light as described previously (27). For conditional mutants, like 8*ΔDGC-1*, in which one of the *alr1219* gene expressions was controlled by the artificial CT promoter (20), cells were grown first in BG11 medium with 0.3 μM copper and 1mM theophylline (TP) to maintain gene expression until logarithmic phase, and then washed three times to avoid inducer effect and transferred to new fresh copper and TP-free BG11 medium to inactivate the gene expression. Neomycin (25 μg/ml), streptomycin (2.5 μg/ml), or spectinomycin (5 μg/ml) was provided in the culture as needed. All strains used in this study are listed in Table S1.

### Suppressor screening and whole-genome sequencing

To isolate suppressors, the conditional mutant 8ΔDGC-1, which exhibits growth arrest under inducer-free conditions, was cultured on copper- and TP-free BG11 plates until new colonies emerged. Briefly, 8*ΔDGC-1* cells grown in BG11 medium supplemented with 0.3 μM copper and 1mM theophylline (TP) were washed three times, serially diluted, and plated onto fresh copper- and TP-free BG11 plates. One month later, newly formed colonies were subjected to primary screening to exclude mutations in the CT promoter and petR (*alr2569*, a copper-dependent promoter inhibitor) (28) by PCR sequencing with primers CT-F/CT-R and *alr2569F/alr2569R*, respectively. Confirmed colonies (no mutations in these genes) were then propagated in fresh copper-free BG11 medium and submitted for whole-genome sequencing (Beina, Wuhan) to identify secondary mutations. All primers used in this study are listed in Table S2.

### Mutant construction and DNA manipulation

The markerless deletion mutants, conditional mutants, and strains with point mutations were created by the genome editing technique based on CRISPR-Cpf1 as previously described (21, 29). To construct the mutant strains, the corresponding plasmid was transferred into *Anabaena* by conjugation as previously described (27). All mutant strains were confirmed by PCR. The primers and plasmids used in this work are listed in Table S2 and Table S3, respectively.

### Extraction and Quantification of Intracellular c-di-GMP

Intracellular c-di-GMP in *Anabaena* was extracted and quantified as previously described (21). In brief, 3OD of cells were collected by filtration when cell density was near OD_750_ = 0.6 and resuspended with 600μL ice-cold extraction buffer (40% acetonitrile, 40% methanol, and 20% ddH_2_O). The cells were broken, incubated ice-cold, and then centrifuged. Then, the supernatants were evaporated and dried by vacuum manifold. c-di-GMP levels were quantified by UPLC-MS/ MS on ACQUITY UPLC H-classXevo TQ MS system equipped with a Waters BEH C18 2.1×50mm column. A standard curve was generated using c-di-GMP (Biolog) at concentrations of 400, 200, 100, 50, 25, 12.5, and 6.25 nM to calculate the c-di-GMP concentration in each sample. All experiments were performed in triplicate.

### Microscopy and Measurement of Cell Size

All microscope images were captured by an SDPTOP EX30 microscope and processed using ImageJ without deconvolution as described (17, 21, 27). For cell size parameter analysis, microscopy images were captured when the optical density of a culture reached 0.4-0.6 at OD750. The cell length and width of 200 cells of each strain were measured by ImageJ. Statistical tests and plotting of data were performed with Origin 8.0.

## Acknowledgments

We would like to thank Jun Men from the Analysis and Testing Center of the Institute of Hydrobiology, Chinese Academy of Sciences, for support in the measurement of c-di-GMP using LC–MS/MS.

## Author contributions

C.Z. and X.Z. designed research; Q.S., Z.D. performed research; Q.S., X.Z. analyzed data; Q.S. X.Z and C.Z. wrote the manuscript.

## Data and materials availability

Data are available in the published article and its online supplemental material.

